# A mechanistic framework for nascent RNA-seq simulations

**DOI:** 10.1101/2025.11.06.687082

**Authors:** Ezequiel Calvo-Roitberg, Jesse W. Lehman, Edric Tam, Shaimae Elhajjajy, Barbara E. Engelhardt, Athma A. Pai

## Abstract

Recent advances in nascent RNA sequencing promise profound insights into transcriptional dynamics. Analyzing the complex datasets generated from nascent RNA genomic approaches require robust computational pipelines and methods to harmonize across experimental workflows. Current solutions include simulation of read counts from individual protocols, but these methods fail to properly replicate biological and technical artifacts or allow for the comparison between nascent RNA approaches. To address this, we introduce *simulated pre-mRNA and RNA kinetics* (*SPARK*), a comprehensive *in silico* framework that generates nucleotide-resolution benchmarking datasets for both nascent short- and long-read sequencing. *SPARK* simulates the entire nascent RNA sequencing process, going from single-molecule transcription and co-transcriptional processing to biochemical enrichment and finally producing high-throughput sequencing reads. By specifically modeling kinetic parameters, such as polymerase speed and processing rates, *SPARK* outputs reads that replicate sources of biological variability alongside technical noise. We demonstrate that *SPARK* effectively benchmarks different existing analytical workflows and evaluate how well indirect read-based metrics reflect underlying biological features. By providing a standard for modeling RNA biogenesis, SPARK can be used to refine mathematical models, design targeted error-correction strategies, and integrate disparate nascent RNA sequencing datasets.

---

RNA sequencing (RNA-seq) has transformed transcriptome analysis by enabling precise and accurate gene expression estimation. However, standard RNA-seq approaches only provide a static snapshot of steady-state mature mRNA levels, which limits our ability to understand the gene regulatory mechanisms that give rise to dynamic mRNA profiles across cellular contexts. Instead, it is increasingly clear that directly characterizing nascent RNA levels and transcriptional dynamics may yield greater insights into the mechanistic steps that underlie mRNA biogenesis.

In the past few years, many approaches have been developed to selectively analyze ongoing or recently completed RNA synthesis. These can be broadly categorized into three groups. First, techniques to assess the precise position of RNA polymerase (RNAP), such as Precision Run-On sequencing (PRO-seq^1^) and Native Elongating Transcript sequencing (NET-seq^2–4^), inherently sequence RNA associated with ongoing transcription. Second, nascent transcripts can be captured by isolating chromatin-associated RNA or RNA associated with other sites of mRNA biogenesis^5^. Finally, the absolute abundance of newly synthesized RNA made in a given amount of time can be quantified with metabolic labeling using nucleotide analogs, such as 4-thio-uridine (4sU^6^) or bromouridine (BrU^7^). Metabolic labeling approaches are increasingly paired with nucleotide recoding chemistry (NR-seq^8,9^), to chemically mark and identify nascent transcripts without the need for selective isolation. Together, these strategies offer a high-resolution, real-time view of transcriptional activity with a temporal resolution that mature RNA sequencing cannot provide. However, since these methods capture nascent RNA during different stages and each have inherent biases, it is hard to know how to properly integrate biological insights gleaned from independent analyses.

The growing variety and adoption of nascent RNA sequencing methods thus comes with a need for robust computational tools that accurately model transcriptional processes. While nascent RNA-specific analytical tools have become more prevalent^9–12^, there is not yet a robust simulation framework to generate realistic benchmarking datasets and validate methods that quantify co-transcriptional processes. Most currently available pipelines simulate nascent RNA^13–16^ by generating aggregated count matrices across nucleotides or genes. They do not simulate sequencing reads, thus, they fail to capture the biological (e.g., variability in kinetic parameters across a gene) and technical biases introduced by library preparation (e.g. edge effects, nucleotide substitutions) and sequencing (e.g. variable read lengths, mapping errors). Furthermore, any read-based simulations are often specific for short- or long-read sequencing methodologies, limiting the evaluation of how well computational or modeling approaches work across them.

To overcome these limitations, we introduce *simulated pre-mRNA and RNA kinetics* (*SPARK*), a comprehensive simulation framework for generating nascent RNA benchmarking datasets (**Fig. 1A**). SPARK uses real annotated gene sequences from any species to simulate nascent RNA sequencing reads after performing *in silico* transcription and processing (**Fig. 1B, left**), nascent RNA enrichment (**Fig. 1B, right, Fig. S2A**), and library preparation steps that include optional nucleotide substitution, sequencing errors, enrichment background, among others (**Fig. S2B**). To optimally capture diverse gene sequences and architectures that may bias experimental or analytical steps, *SPARK* has an option to first cluster genes based on several relevant features and chooses a user-determined number of genes across clusters (**Fig. S1A, Methods**).

**Figure 1.**
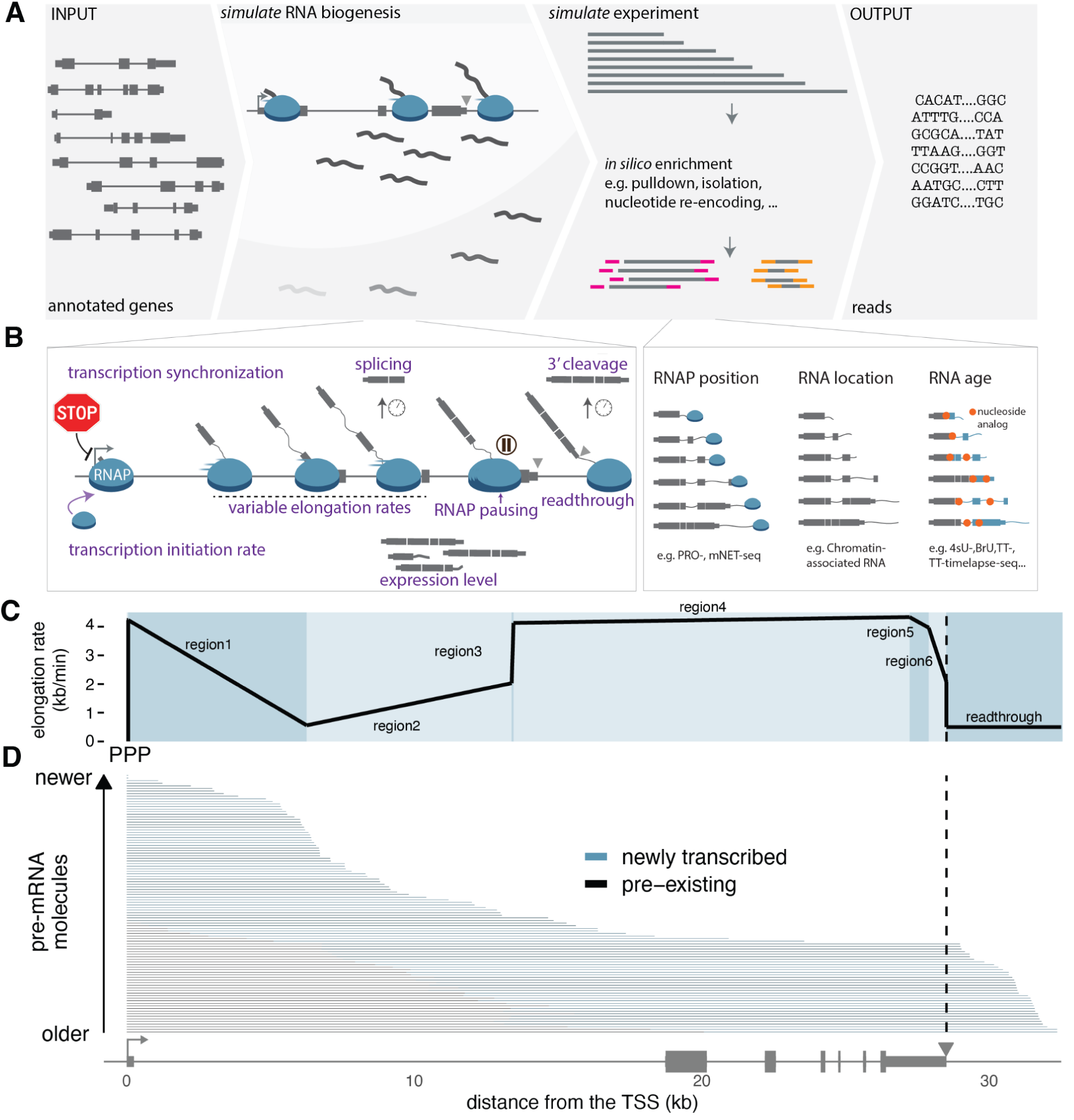
*SPARK* simulates reads from nascent RNA sequencing methods. **(A)** Schematic overview of *SPARK* **(B)** *SPARK* allows for the modulation of various biological parameters (*left*) to generate nascent transcripts prior to the choice of *in silico* nascent RNA enrichment strategies (right). (**C**) Simulated elongation rate variation across the *BAHD1* gene, with a promoter-proximal pause (PPP) and 7 distinct elongation blocks. (**D**) Simulated pre-RNAs from *BAHD1* shown as horizontal lines, with pre-existing (mature; *black*) and newly-transcribed (nascent; *blue*) regions delineated for each molecule.

## Simulation of RNA transcription

Previous studies have shown that RNAP elongation rates are highly dynamic both within and across genes^17–19^. To capture this complexity, SPARK can simulate both uniform and variable elongation kinetics by assigning nucleotide-specific RNAP speeds along the entire gene, extending past the polyadenylation site (PAS) into the downstream readthrough region by defining elongation rate blocks (**Methods; Fig. 1B, left panel; Fig. S1B**). Within each block, the elongation rate is a linear interpolation between the first and last nucleotide of the region. This allows for the simulation of continuous acceleration, deceleration, or constant elongation at single-nucleotide resolution, reflecting the variable kinetics of elongating RNAP. Furthermore, *SPARK* allows for the modulation of transcription initiation rates, the incorporation of tunable promoter-proximal (PPP) or gene-body RNAPII pausing events, and the simulation of time-course assays involving the pharmacological synchronization of transcription (e.g., 4sUDRB-seq) (**Fig. 1B**).

As a representative example, we simulated nascent RNA reads across assays from the *BAHD1* gene, including an initial PPP, six elongation rate blocks across the gene body, and a final post-PAS readthrough region (**Fig. 1C**). Based on chosen kinetic parameters, *SPARK* performs *in silico* synthesis of each individual RNA molecule over experimental time. The framework classifies transcript segments into “pre-existing” (synthesized prior to the simulated experiment) and “newly transcribed” (synthesized during the simulated experiment) (**Fig. 1D**). Consequently, RNAP that initiate transcription within the experimental time produce RNA composed entirely of newly transcribed sequences. Furthermore, because *SPARK* tracks RNAP position over time, regions with pausing or slower elongation accumulate higher densities of nascent RNA. This recapitulates the *in vivo* accumulation of RNAP observed at bottleneck regions in nascent transcriptomics data^1,2,20^.

## Simulating RNA processing

RNA processing events, such as splicing and 3’ end cleavage, can occur either co-transcriptionally, while RNAP is actively elongating, or post-transcriptionally. To capture these temporal dynamics, *SPARK* simulates the kinetics of both processes at the single-molecule level. Briefly, splicing and cleavage are simulated probabilistically based on a first-order exponential decay distribution, as described previously^21^.

Splicing kinetics are driven by an intron-specific half-life, defined as the time since synthesis of the 3’ splice site (3’ss) at which 50% of the transcript population has successfully undergone splicing (**Fig. 2A**). To illustrate this, we simulated the transcription and processing of the *PRKAR1B* gene. At the single-molecule level, older transcripts have a higher probability of being spliced due to the older lifetime of their respective 3’ss (**Fig. 2B**). Across the population of simulated transcripts, splicing is exponentially associated with the time since 3’ss synthesis such that 50% of molecules have been spliced at the defined intron half-life of 3.42min (**Fig. 2C**). To further confirm the simulation of splicing rates at a larger scale, we used SPARK to generate 4sU-seq datasets simulating 5, 10, and 15 minutes of metabolic labeling. SPARK accurately simulates intron removal within transcripts based on the assigned half-lives (**Fig. S3A**). Using an published approach to estimate intron half-lives from empirical 4sU-seq data^21^, we see that intron half-lives estimated from *SPARK* are highly correlated with the input half-lives (Pearson’s *r* = 0.89) (**Fig. S3B**).

**Figure 2.**
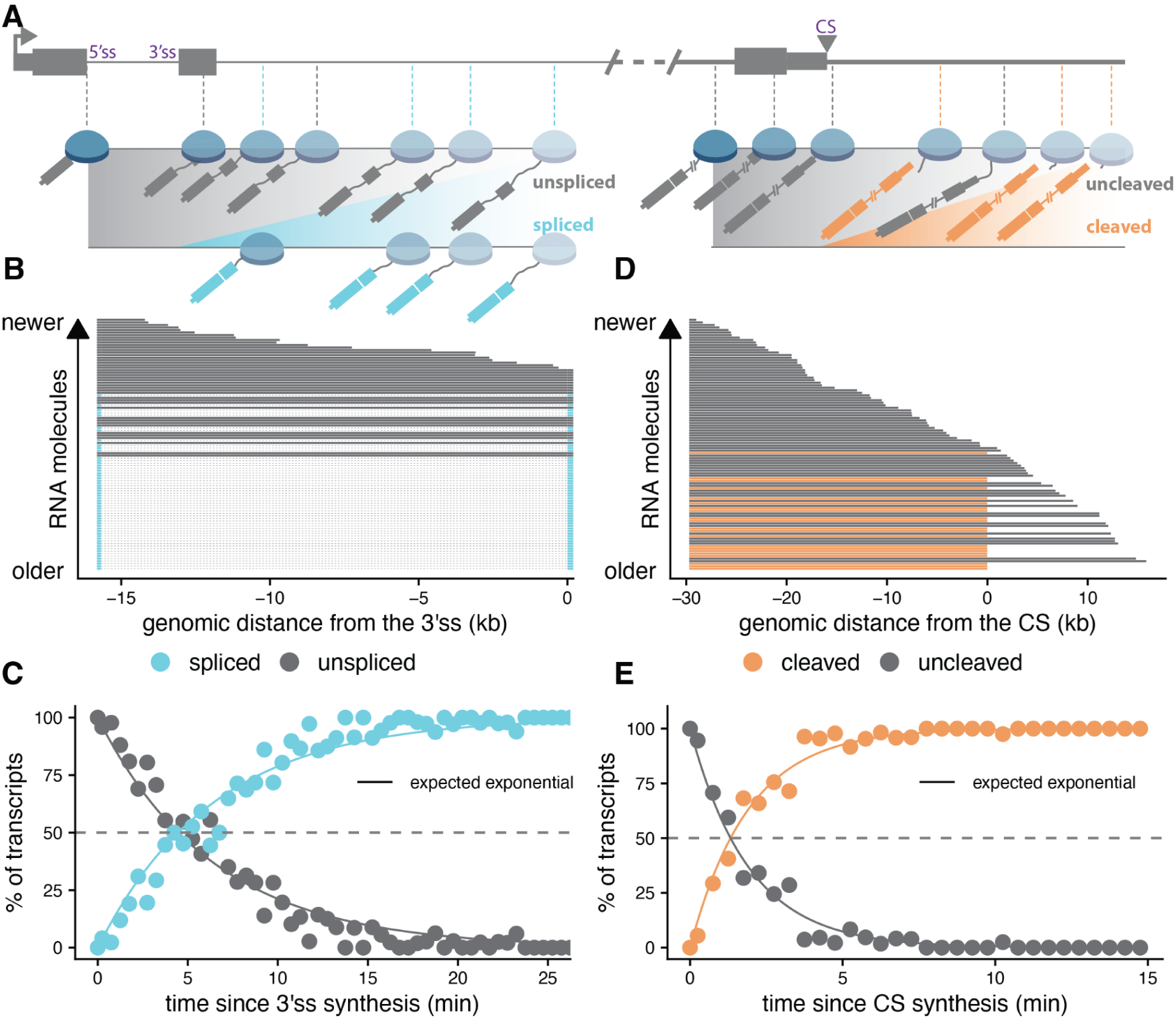
*SPARK* simulates RNA processing kinetics. (**A**) Conceptual representation of probabilistic splicing and cleavage kinetics in the simulations, where the probability of sampling reads from processed transcripts is dependent on intron- or cleavage-site (CS) specific half-lives and location of the polymerase at the completion of the experiment. (**B**) Simulated transcripts for the *PRKAR1B* gene, zooming in the region around the first two introns and where older molecules are more likely to be spliced. (**C**) Proportion of unspliced (*grey*) or spliced (*blue*) transcripts relative to time since 3’ splice site (ss) synthesis. The distribution is the expected exponential increase or decrease for the simulated intron half-life of 4.16min. (**D**) Simulated transcripts for the *SMIM7* gene, where older molecules are more likely to be cleaved. (**E**) Proportion of uncleaved (*grey*) or cleaved (*orange*) transcripts for the SMIM7 gene relative to the time since cleavage site synthesis. The distribution is the expected exponential increase or decrease for the simulated cleavage half-life of 1.27min.

We employed a similar probabilistic approach to simulate 3’ end cleavage. For each cleavage site (CS), the simulation uses a cleavage half-life, defined as the time at which 50% of the molecules that have been elongated past the CS are cleaved (**Fig. 2A**). We simulated the 3’ end processing dynamics for the gene *SMIM7*. We see that, analogous to splicing rates, (1) older molecules show a higher likelihood of being cleaved due to the older lifetime of their CS (**Fig. 2D**) and (2) cleavage is exponentially associated with the time since CS synthesis such that 50% of transcripts are cleaved at the simulated CS half-life for this gene (**Fig. 2F**). Finally, we again see that the expected and the observed proportion of cleaved molecules from simulations are highly correlated (**Fig. S3C**). Together, these tests demonstrate that *SPARK* accurately simulates splicing and 3’ end cleavage to produce synthetic reads suitable for benchmarking mathematical approaches to model RNA processing kinetics from metabolic labeling data.

In addition to modeling 3’ end cleavage kinetics and RNAP elongation rates in the readthrough region, SPARK simulates the degradation of downstream RNAP-associated cleavage intermediates. This readthrough degradation (Rdeg) is again modeled with a first order exponential decay rate (**Fig. S3D**). To test the dynamic range of this module, we simulated a 10 minute metabolic labeling experiment and calculated the readthrough index (RTI) across simulated Rdeg values. The simulated read density for the *SF3B4* gene shows a progressive decrease in coverage downstream of the CS with increasing Rdeg (**Fig. S3E**). Extended to a larger scale, the RTI steadily decreases as the Rdeg increases, with the RTI going to 0 when Rdeg equals the RNAP elongation rate, at which the nascent readthrough transcript is expected to be degraded as fast as it is synthesized (**Fig. S3F**).

## *In silico* nascent RNA enrichment and library preparation

Following the generation of transcripts using the kinetic framework described above, *SPARK* can simulate reads from a wide array of nascent and mature RNA sequencing assays. Users can choose modules to simulate model RNAP position, chromatin retention, time-resolved metabolic labeling, or steady-state mature RNA. For metabolic labeling techniques like 4sU-seq or TT-seq, *SPARK* models the stochastic incorporation of nucleotide analogues (e.g., 4-thiouridine, 5-ethynyluridine, 2-bromouridine, 6-thioguanine). For metabolic labeling modes, users can also select nucleoside recoding to induce substitutions at labeled positions (e.g. T > C substitutions like those induced for 4sU-containing positions^8,9,21^). To recreate run-on assays like PRO-seq, *SPARK* simulates transcription termination driven by biotin-NTP (**Fig. S2A**). By specifying the total time available for *in silico* RNA synthesis, users can set the time available for all kinetic parameters to occur (e.g. for how long RNAP can elongate).

Following transcript synthesis, *SPARK* mirrors molecular enrichment steps for the chosen experimental protocol. This includes simulating pulldown-based enrichments for nucleotide analogs or RNAP (with optional fragmentation for NET-seq) and biochemical fractionation for chromatin-associated RNA. To simulate this spatial fractionation, *SPARK* includes a chromatin export half-life parameter. This metric defines the time at which 50% of the fully cleaved transcripts for a given gene are released from the chromatin. Uncleaved nascent transcripts and downstream RNAP-associated cleavage products remain chromatin-associated by default. However, fully cleaved RNAs are partitioned between the chromatin and nucleoplasmic fractions strictly based on the time elapsed since their individual cleavage events. Using this parameter, we see that the simulation proportion of chromatin-associated molecules is strongly correlated (Pearson’s r = 0.97) with the expected proportion based on theoretical distributions such that transcripts with shorter export half-lives have a higher fraction of non-chromatin-associated molecules (**Fig. S3G**).

To illustrate the versatility of *SPARK*, we simulated reads from six distinct experimental protocols using an identical set of kinetic parameters and simulated RNAs for the *RALGDS* gene. The simulation was conducted with a PPP event followed by eight regions of varying RNAP elongation rates (**Fig. 3A bottom track**). After each *in-silico* enrichment, short reads were generated and mapped. The resulting coverage recapitulates characteristic features of empirical data, with expected distributions and biases. For instance, simulated reads from methods designed to map actively engaged RNAP positions, such as PRO-seq and NET-seq, are enriched at the 5’ end of the gene, with a sharp peak corresponding to the simulated PPP (**Fig. 3A**). In contrast, methods that capture broader temporal windows, including metabolic labeling (4sU-seq and TT-seq) and chromatin-associated RNA fractionation, result in a wider distribution of read coverage across the entire gene body. As expected, the simulated TT-seq profile demonstrates less 5’ bias relative to the 4sU-seq profile, reflecting the known biochemical biases of these specific enrichment strategies. Finally, the simulated steady-state mature RNA has read coverage exclusively over exonic regions. By generating assay-specific reads from an identical set of simulated biological parameters, *SPARK* provides controlled datasets to directly compare and calibrate analytical pipelines across distinct sequencing methods.

**Figure 3.**
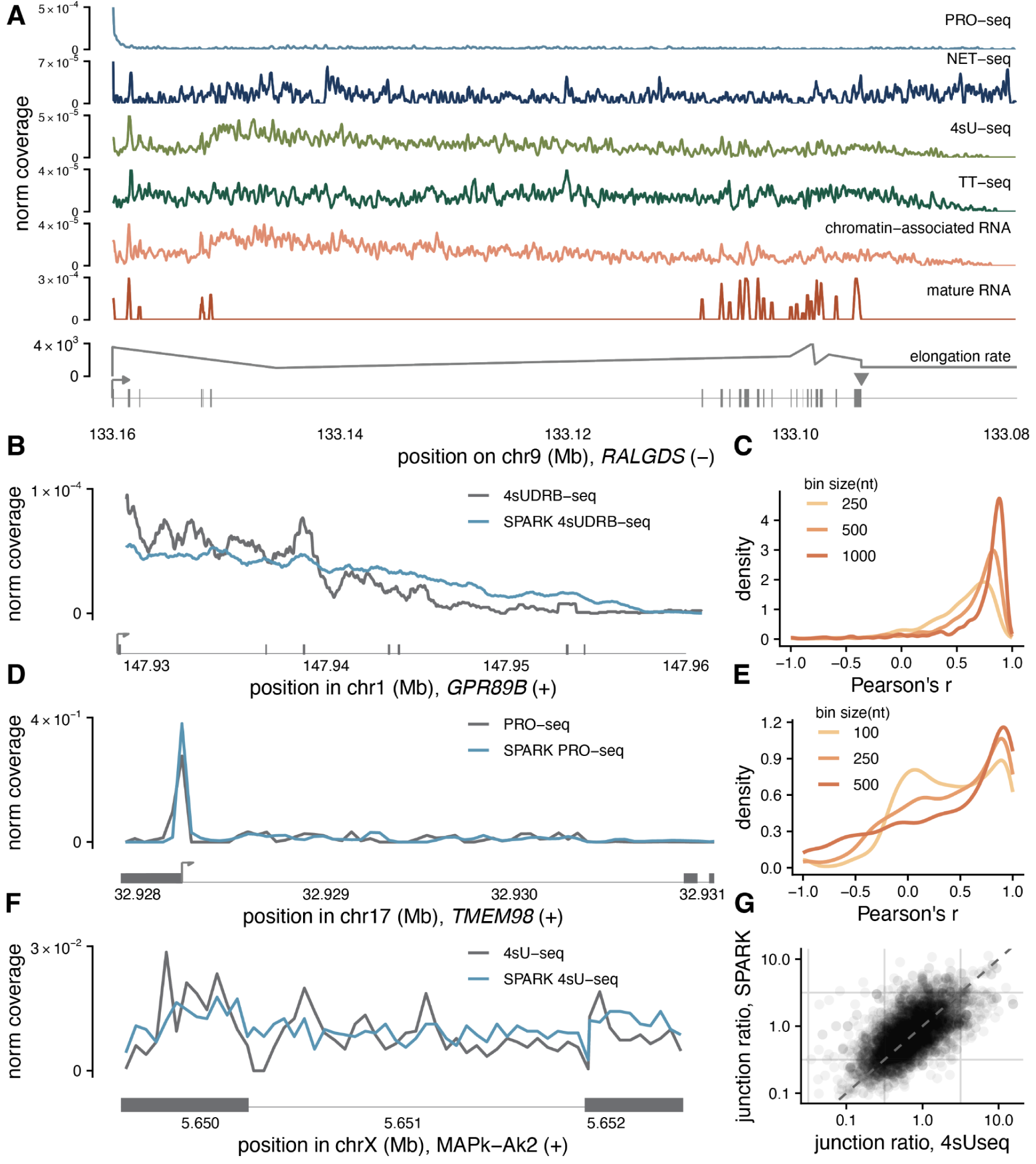
*SPARK* simulates and replicates nascent RNA experimental protocols. (**A**) Comparison of normalized read coverage across six nascent RNA approaches (PRO-seq, NET-seq, 4sU-seq, TT-seq, chromatin-associated RNA, and mature polyadenylated RNA) simulated with the same biogenesis kinetics parameters for the *RALGDS* gene. The bottom tracks display the assigned elongation rates per nucleotide, with an initial promoter-proximal pause followed by eight distinct elongation rate blocks and the gene structure (TSS indicated with an arrow, exons indicated as boxes, *bottom*). (**B**) Comparison of empirical (*grey*) and SPARK-simulated (*blue*) 4sUDRB-seq coverage across the human *GPR89B* gene. (**C**) Distribution of Pearson correlation coefficients between empirical and simulated 4sUDRB-seq coverage across genomic bin sizes of 250, 500, and 1000 nt (n = 1,474 genes). (**D**) Comparison of empirical (*grey*) and *SPARK*-simulated (*blue*) PRO-seq coverage across the human *TMEM98* gene. (**E**) Distribution of Pearson correlation coefficients between empirical and simulated PRO-seq coverage in the promoter region across genomic bin sizes of 100, 250, and 500 nt (n = 4,481 genes). (**F**) Comparison of empirical (*grey*) and *SPARK*-simulated (*blue*) 4sU-seq coverage profiles across the fly MAPk-Ak2 gene. (**G**) Genome-wide comparison between the ratio of unspliced to spliced junction reads from experimental (*x-axis*) and *SPARK*-simulated (*y-axis*) reads generated from *Drosophila melanogaster* 4sU-seq datasets (n = 17,670 introns).

We next wanted to confirm that *SPARK* accurately simulates the technical parameters associated with high-throughput sequencing library preparation. We looked at how well metrics estimated from simulated reads recapitulate the simulated ground truth for three different features: gene expression levels, insert sizes generated from fragmentation and size selection during short-read RNA-seq library preparation, and sequencing error rates (**Fig. S4A-C**). For all three of these parameters, the estimates from simulated reads are highly correlated with the ground truth (Spearman or Pearson correlations > 0.9 for all parameters).

## *SPARK* accurately replicates empirical data characteristics

To evaluate whether *SPARK* accurately reproduces technical and biological features of empirical nascent RNA datasets, we used published kinetic parameters derived from specific sequencing approaches to simulate reads from the corresponding approach. We note that since most studies typically measure only a single kinetic parameter, we set unmeasured parameters to be constant in the simulations, which may account for some variance relative to experimental data. Importantly, this demonstrates *SPARK’s* utility as an *in silico* framework where confounding biological variables can be isolated; by holding sets of parameters constant, the framework enables the precise identification of how individual parameters uniquely shape sequencing outputs.

First, we evaluated if *SPARK* could accurately recapitulate human gene-specific RNAPII elongation rates derived from 4sUDRB sequencing data^18^. In this assay, DRB treatment is used to stall RNAPII after transcription initiation (forestalling transition into active elongation), followed by release and labeling with 4sU for a short time. The resulting 4sU labeled nascent RNA is derived from a synchronized population of RNAPII molecules, allowing the estimation of how far RNAPII can travel in a given time defined by the labeling period. We simulated 4sUDRB-seq reads with *SPARK*, using gene-specific elongation rates estimated by Fuchs *et al.* 2014. We see that the distribution of simulated reads for the *GPR89B* gene mirrors coverage from the empirical 4sUDRB-seq dataset and captures the position of the advancing RNAPII wavefront (**Fig. 3B**). Coverage profiles across genes were also highly correlated between simulated and experimental data (**Fig. 3C**).

Next, we evaluated if *SPARK* could accurately recapitulate read profiles derived from approaches to estimate RNAP position. We used *SPARK* to simulate PRO-seq, a nuclear run-on assay that incorporates biotinylated NTPs to terminate nascent transcript elongation and enable single-nucleotide resolution mapping of active transcription^1^. We simulated reads using PPP durations empirically estimated from time-course PRO-seq for 2, 8, and 32 minutes of run-on ^22^. The distribution of simulated PRO-seq reads for the *TMEM98* gene (**Fig. 3D**) was similar between simulated and experimental datasets, particularly within promoter-proximal regions. More broadly, the coverage at promoter regions was highly correlated between simulated and empirical datasets (**Fig. 3E**).

Finally, we evaluated if *SPARK* could accurately recapitulate read profiles around exons and introns that reflect variable splicing dynamics. We simulated fly transcripts using empirically derived *Drosophila melanogaster* intron half-lives estimated from 4sU-seq data^21^, where cells are incubated with 4sU for short pulses and 4sU-labeled RNA is selectively isolated for sequencing. The distribution of simulated 4sU-seq reads around a given intron in the MAPk−Ak2 gene (**Fig. 3F**) shows similar coverage between simulated and empirical data. We used a ratio of intron-exon to exon-exon junctions to summarize the splicing profile for a given intron and see that this ratio is highly correlated across introns between simulated and empirical data (**Fig. 3G**). Collectively, these benchmarking experiments demonstrate that SPARK accurately simulates reads that replicate empirical nascent RNA sequencing profiles across diverse experimental settings, regulatory mechanisms, and temporal scales.

## *SPARK* enables benchmarking of nascent RNA analytical frameworks

*SPARK* provides a unified computational framework to benchmark diverse nascent RNA sequencing workflows across both short- and long-read technologies. Because *SPARK* generates nucleotide-resolution reads rather than matrix counts, it enables the evaluation of tools that rely on read-level features such as nucleotide recoding events or precise fragment ends to estimate transcriptional dynamics. Furthermore, *SPARK* allows us to evaluate how well indirect, read-based metrics perform at approximating direct biological parameters under varying technical biases. To demonstrate this, we used *SPARK*-simulated reads to evaluate three established computational workflows.

First, we used *SPARK* to benchmark GRAND-SLAM^10^, a package designed to analyze SLAM-seq data ^8^. SLAM-seq is a nucleoside recoding approach where 4sU labeled RNA is treated with iodoacetamide to induce T-to-C conversions at 4sU-containing nucleosides during first-strand synthesis. After sequencing of the total RNA population, these conversions are used to distinguish reads derived from nascent transcripts relative to reads from pre-existing RNA. To quantify the proportion of nascent molecules for each gene, GRAND-SLAM uses probabilistic modeling to identify converted reads and calculate a new-to-total ratio (NTR). To evaluate the accuracy of this estimation, we used *SPARK* to simulate reads from an eSLAM-seq experiment (where the “e” stands for 4sU enrichment prior to sequencing^23^), with a range of background total RNA molecules without conversions to represent varying pre-existing RNA contamination in the dataset. GRAND-SLAM estimates are broadly correlated with simulated NTRs across varying proportions of background (non-nascent) RNA (**Fig. 4B**; Pearson’s r = 0.85). However, this accuracy is dependent on the probability of T-to-C conversions, which scales with the extent of 4sU incorporation or labeling. We observe that GRAND-SLAM NTR estimates are more accurate at longer simulated 4sU labeling times (**Fig. S4D**), which increases the amount of labeled RNA. This is an important consideration for the design of nucleotide recoding experiments, since labeling times can range from a few minutes to hours. We reasoned that the underestimation of NTR under certain conditions stems from GRAND-SLAM’s estimation of 4sU incorporation rates. To test this, we used *SPARK* to generate SLAM-seq datasets with varying incorporation rates (defined as the probability of a U to be a 4sU). We see that GRAND-SLAM consistently underestimates the ground truth *in silico* incorporation rate (**Fig. S4E**) and this underestimation is more pronounced as the incorporation rate increases, likely driven by mapping errors.

**Figure 4.**
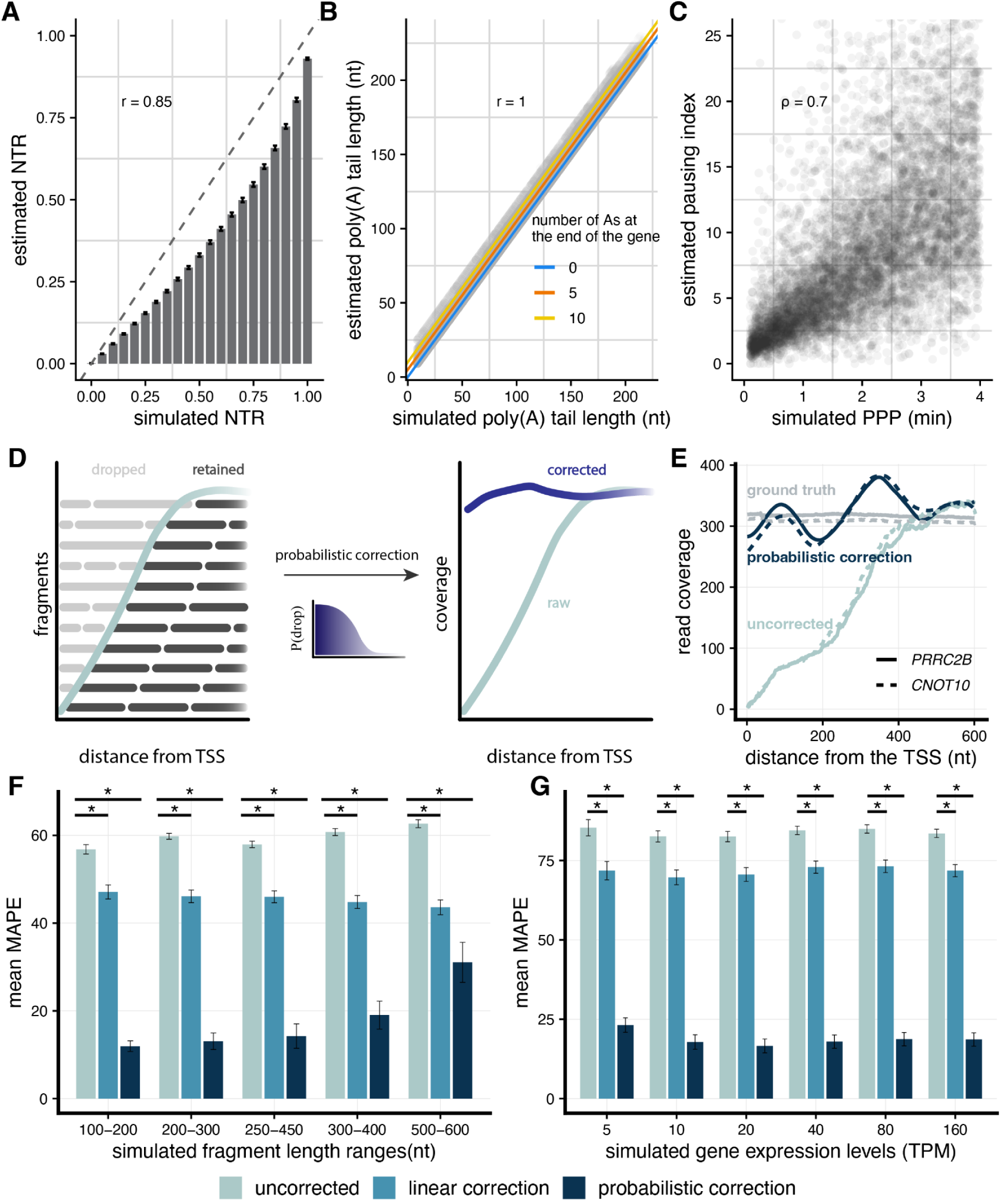
*SPARK* enables benchmarking and optimization of RNA sequencing analysis pipelines. **(A)** Benchmarking the estimation of the new-to-total ratio (NTR) from GRAND-SLAM. Each bar represents the mean estimated NTR (*y-axis*) across simulated NTR values (*x-axis*), where error bars are standard error of the mean (n = 1,000 genes per bar) and the dashed line represents y=x line. **(B)** Correlation between simulated poly(A) tail lengths (*x-axis*) and lengths estimated by the FLAM-seq pipeline (*y-axis*) (n=200 genes). The lines represent linear fit for transcripts with 0 (blue), 5 (orange), or 10 (yellow) templated As at the end of the read. **(C)** Relationship between the simulated promoter-proximal pausing (PPP) duration (*x-axis*) and the Pausing Index (PI) calculated from simulated PRO-seq reads (*y-axis*, n = 8000 human genes). **(D)** Schematic of 5’ edge effect correction. The edge effect is characterized by fragment dropout near the TSS during fragmentation and size selection. **(E)** Gene coverage for *PRRC2B* (*solid lines*) and *CNOT10* (*dashed lines*) showing raw uncorrected coverage (*light blue*), probabilistically corrected coverage (*dark blue*), and ground-truth signal before fragmentation and size selection (*grey*) as a function of distance from the TSS. **(F)** Comparison of mean absolute percentage error (MAPE) across varying simulated fragment length ranges (nt) for uncorrected (*light blue*), linear corrected (*medium blue*), and probabilistic corrected (*dark blue*) data. Error bars represent standard error. * adjusted p value<10^-9^. **(G)** Comparison of mean MAPE across a range of simulated transcript per million (TPM) uncorrected (*light blue*), linear corrected (*medium blue*), and probabilistic corrected (*dark blue*) data. Error bars represent standard error. *adjusted p value<10^-9^.

Second, we evaluated the FLAM-seq analysis pipeline^24^, designed to quantify poly(A) tail lengths from FLAM-seq long reads. We generated PacBio-style long reads from transcripts with varying simulated poly(A) tail lengths and ran the FLAM-seq pipeline. Poly(A) tail lengths estimated from simulated reads mostly concur with ground-truth values (Pearson’s r = 1; **Fig. 4A**). However, we see that this pipeline can overestimate poly(A) tail lengths for transcripts ending in templated adenosines, with a skew that is dependent on the number of templated terminal adenosines.

Third, we used *SPARK* to benchmark metrics developed to infer kinetic parameters from genomic approaches that lack temporal resolution. Specifically, we evaluated the Pausing Index (PI), an indirect measurement that quantifies RNAPII PPP using the ratio of reads within the promoter-proximal region relative to the gene body^25–27^. By varying the PPP durations in *SPARK*, we tested how accurately this indirect read-based metric reflects true pausing kinetics. We simulated PRO-seq reads with varying PPP durations and observed a strong positive correlation between the PI and simulated pause duration (**Fig. 4C**, Spearman’s ρ=0.7), confirming that the PI generally approximates temporal dynamics of pausing, particularly for events under two minutes. The observed linear relationship between these metrics suggests that a conversion factor could be developed to translate empirically estimated PIs into absolute pausing durations.

Finally, to demonstrate the utility of *SPARK* to benchmark new analytical approaches, we used it to validate and fine-tune a novel computational method we developed to correct RNA-seq edge effects. During short-read library preparation, random fragmentation followed by size selection leads to a depletion of fragments, and ultimately sequencing coverage, at the ends of RNA molecules. This process leaves these regions systematically underrepresented^28–30^ in RNA-seq data. This limitation is particularly problematic when mapping RNAP positions or analyzing nascent RNA data, as both workflows derive substantial biological insights from the 5’ regions of mRNA molecules. Using *SPARK*-simulated reads to evaluate the edge effect within the 5’ regions of genes, we developed a correction approach that models the base-specific probability of fragment dropout during size selection (**Fig. 4D-E**). These estimated dropout probabilities are used to calculate the expected coverage loss at each base, which in turn informs the correction for the resulting bias (Methods). While a linear correction improves edge coverage relative to uncorrected data (average 13.2% decreased MAPE), our probabilistic correction provides a greater improvement across a range of fragment lengths (average 41.7% decreased MAPE; **Fig. 4F**) and gene expression levels (average 65.1% decreased MAPE; **Fig. 4G**). This shows that *SPARK* can be used to develop new computational analytical approaches and fine tune data correction algorithms.

## Conclusion

The expansion of nascent RNA sequencing technologies has provided an unprecedented view of transcriptional regulation. Yet, reconciling data across these methods presents a significant analytical hurdle, as each protocol relies on specific biochemical enrichments with distinct targets and properties that inherently bias the snapshot of the RNA lifecycle they capture. Consequently, integrating these disparate datasets remains difficult. Furthermore, the reliance on count-based simulations as ground truth limits the ability to rigorously evaluate how technical biases introduced during nascent RNA enrichment, library preparation, sequencing, and read mapping impact downstream analyses. *SPARK* bridges this gap by generating nucleotide-resolution short or long reads across a large diversity of nascent RNA genomics approaches, fully simulating the biological and experimental steps from transcription initiation to high-throughput sequencing.

Reads simulated with *SPARK* can help to benchmark existing or novel computational workflows. *SPARK* enables the systematic evaluation of biases related to mechanistic cellular dynamics over time, genomic sequence features, and experimental efficiency. This allows not only the development of well calibrated bioinformatic or mathematical modeling strategies, but also assessment of the lower and upper bounds within which measurements are accurate. Furthermore, these simulations can serve as a tool for evaluating how indirect analytical measurements behave under defined biological parameters. Because some metrics require inferring kinetic parameters from static read distributions, separating true biological signal from experimental and technical noise is often challenging. For instance, we show that the pausing index is highly correlated with simulated pausing durations, validating the use of this metric as an inference for promoter proximal pausing dynamics. Together, these benchmarking efforts allow for a definitive assessment of how accurately established metrics reflect the biological mechanisms they are designed to capture and measure.

Ultimately, *SPARK* provides a unified computational framework to correct and integrate data across diverse sequencing methods. As transcriptomic analyses become increasingly complex, the ability to translate read profiles into a cohesive model of RNA biogenesis is paramount. By modeling the biological and technical parameters that give rise to assay-specific biases, *SPARK* facilitates the development of correction algorithms, harmonizing seemingly conflicting datasets. This framework not only standardizes the development and benchmarking of current analytical tools but also establishes a scalable, predictive foundation to support the next generation of nascent RNA sequencing methods.

## CODE AVAILABILITY

*SPARK* and the code for downstream analyses included in this manuscript are available at https://github.com/thepailab/spark. The edge effect corrector implemented in this manuscript is available at https://github.com/edrictam/EdgeEffectCorrector.

## ACKNOWLEDGEMENTS

We thank Alexander Park for work on early versions of the simulation scripts and Adam Siepel, Engelhardt lab, and Pai lab for helpful discussions, suggestions, and comments, particularly to Eva Jazbec, Valeria Sanabria, and Kelly Cochran. This work was funded by grants R35GM133762 (AAP), R01HG012944 (AAP), R01HG013736 (BEE), and R01HG012967 (AAP and BEE) from the National Institutes of Health, the Parker Institute for Cancer Immunology (PICI), the Biswas Family Foundation, the Warren Alpert Foundation (ET), the Croucher Foundation (ET) and the Chan-Zuckerberg Institute (BEE). BEE is a CIFAR Fellow in the Multiscale Human Program.

## METHODS

### *SPARK* framework

The basic *SPARK* framework involves (1) selection of genes, (2) assignment of elongation rates across a gene, (3) *in silico* simulation of RNA molecule transcription and processing, and (4) simulating enrichment of nascent RNA through one of five experimental modes, with many optional parameters to modify experimental conditions as desired.

#### Gene clustering and selection

If specified, *SPARK* can select genes from which to simulate nascent RNA transcripts that represent a diversity of genomic features. Users specify the number of genes from which they would like to simulate reads and provide: (1) a gtf with gene annotations and (2) a reference fasta file. For each gene, the longest isoform (upstream-most transcript start site and downstream-most transcript end site) is selected, and its features are used for classification. It is possible to simulate multiple isoforms per gene by delineating them as different genes in the reference. Genes in the gtf are first categorized by gene length, total transcript length(s), number of exons, mean exon lengths, number of introns, mean intron lengths, first intron length, lengths of 5′ and 3′ untranslated regions (UTRs), and exonic and intronic nucleotide composition. Hierarchical clustering is used to identify clusters of transcripts with similar characteristics. An equal number of genes is selected from each cluster for downstream simulations, ensuring a balanced and unbiased set of genes reflecting genome diversity. *SPARK* can simulate genes for any species with an annotated genome and also use only selected genes from a pre-filtered gtf file. By default, all genes listed in the gtf will be simulated.

#### Nucleotide-specific elongation rates

*SPARK* simulates variable transcription elongations by assigning a specific rate to each nucleotide within a gene. The user first defines ranges of (a) the number of elongation blocks, (b) the length of elongation blocks, and (c) elongation rates, which *SPARK* uses to assign elongation blocks for each gene. Nucleotides within each block are assigned a specific rate based on a linear interpolation between the rate at the beginning of the block and the rate at the end of the block, assuming a constant change across the entire block. Users can also specify simpler elongation rate profiles, such as a single, constant elongation rate for all nucleotides within a defined block or simulating a constant rate across the gene.

Finally, users can choose to include RNAP pausing events throughout the gene. Pause sites can be placed randomly across the gene or at specific genomic locations (e.g. the TSS). These events are defined as single-nucleotide blocks. The duration of the pause can be user-defined and is used to define the time required to traverse that specific nucleotide. Therefore, a pausing event is equivalent to the RNAP traversing a single nucleotide extremely slowly.

#### Simulation of RNA synthesis

RNA molecules are synthesized *in silico* based on a series of rates that determine the distribution of RNAP positions across a gene at the start, during, and at the end of each simulated experiment. *SPARK* simulates individual initiation events as a Poisson process with a gene-specific initiation rate *i*, where the time interval between consecutive initiation events is drawn from an exponential distribution with mean 1/*i*¹⁷. For each gene, 1/*i* is drawn log-uniformly from a user-defined range. Because the number of molecules a gene produces scales with *i*, this range sets the distribution of expression levels across genes. The total time required for an RNAP to traverse the gene, *T*, is the sum of the traversal times of its nucleotides, 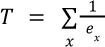 , where *eₓ* is the elongation rate assigned to nucleotide *x*; for pause sites, the traversal time is the pause duration. The average number of polymerases simultaneously transcribing a gene at steady state is therefore 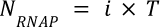

To establish the initial state of the simulation, *SPARK* first populates the gene with *N_RNAP_* that would be present assuming steady-state ongoing transcription. The position of each RNAP is assigned probabilistically. First, RNAPs are assigned to genomic regions (e.g., elongation blocks or pause sites) probabilistically given the relative time required to traverse that region. Second, RNAPs are assigned to precise nucleotide positions chosen from a uniform distribution across the given region. This weighted assignment of RNAP locations ensures that regions with longer traversal times have more RNAP at steady-state.

*SPARK* simulates a target number of total mRNA molecules *N_RNA_* (default 5,000) by calculating the number of independent gene copies *N_gene_* (analogous to DNA molecules across a population of cells) required to generate that number of molecules during the simulation. This is determined by dividing *N_RNA_* by the total mRNAs produced per copy, which is the sum of newly initiated polymerases (*i* × *t*, where *t* is the simulation time) and *N_RNAP_*:

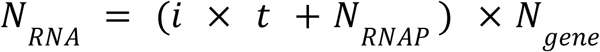

Each gene copy is simulated as an independent transcriptional locus.

#### Sequencing depth and gene expression

The number of reads per gene *SPARK* generates is a consequence of its simulated kinetics. After the experimental mode and library preparation steps, *SPARK* counts the sequenceable units from each gene (e.g. size-selected fragments, NET-seq footprints, run-on fragments, or whole molecules for long reads) and divides by N_gene_ to obtain the units produced per gene copy. The user-defined sequencing depth is distributed across genes in proportion to this per-copy value. As a result, TT-seq read counts scale with *i* × *L*, (where *L* is gene length), NET-seq read counts scale with N_RNAP_, and mature RNA read counts scale with *i* × *length_mRNA_*(where *length_mRNA_* is the length of the mature transcript length. Ground-truth RPKM and TPM values are computed from the reads assigned to each gene.

#### Simulation of RNA splicing

To model the removal of introns from newly synthesized transcripts, *SPARK* simulates RNA splicing as a stochastic, first-order kinetic process, as described previously ^21^. Users define splicing kinetics by providing either intron-specific half-lives or a range of global splicing rates that the framework assigns at random to each intron. During *in silico* synthesis of each RNA molecule, *SPARK* marks an intron as available for excision only once the 3′ splice site (the final base of the intron) has been transcribed. For each intron, SPARK computes the time elapsed between synthesis of the 3′ splice site and the end of the simulated experiment, and models excision as a single-exponential decay, such that the probability that the intron has been removed is:

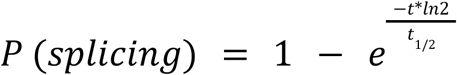

where *t* is the elapsed time since the 3’ss was available and t_1/2_ the assigned half-life. *SPARK* assigns the splicing state of each intron in each molecule using this probability. Introns that are unspliced retain their intronic sequence within the nascent RNA molecule, whereas the sequence of spliced introns is removed and flanking exons are ligated in the simulated transcript.

#### Simulation of RNA cleavage

To model the 3′ end processing and termination of newly synthesized transcripts, SPARK simulates 3′ end cleavage as a stochastic, first-order kinetic process at the cleavage site (CS), as described previously ^28^. Users define cleavage kinetics by providing either a gene-specific cleavage half-life or a global range from which the framework samples a single half-life per gene. During the *in silico* synthesis of each RNA molecule, cleavage can only occur once the RNAP has elongated past the CS. For each transcript, SPARK draws a cleavage time from an exponential distribution defined by the assigned half-life (as described for splicing above) and compares it against the time elapsed between synthesis of the CS and the end of the experiment. If the simulated experiment ends before the cleavage time, the simulated transcript is uncleaved and retains the sequence of the extended readthrough sequence. If the cleavage time elapses before the simulation concludes, *SPARK* cleaves the simulated transcript at the CS, can append a poly(A) tail to the resulting molecule, and treats the downstream readthrough fragment as an independent species that is progressively degraded from the 5’ end with a defined readthrough degradation rate.

### *SPARK* experimental modes

Following the synthesis of RNA molecules, SPARK can generate reads from one of five nascent RNA experimental approaches: metabolic labeling with full-molecule nascent RNA enrichment, metabolic labeling with fragmented nascent RNA enrichment, RNAP run-on, nascent RNA associated with elongating RNAPs, or biochemical fractionation of chromatin-associated RNA.

#### Metabolic labeling and nascent RNA enrichment

*SPARK* simulates metabolic labeling experiments (e.g., 4sU-seq, TT-seq) by first simulating the stochastic incorporation of nucleotide analogs (e.g. 4sU). Users specify the incorporation rate, which defines the probability that an analog will be incorporated instead of the canonical nucleotide (e.g., uridine). The subsequent *in silico* enrichment step is performed differently depending on the specific protocol being simulated (specified below).

#### Nascent RNA enrichment

The nascent RNA enrichment experimental mode simulates a 4sU-seq style nascent RNA pulldown experiment by conditioning on full-length molecules that have at least one nucleotide analog incorporated. Following the simulation of full-length RNA molecules, discards simulated molecules that do not have at least one incorporated nucleotide analog. The remaining set of full-length labeled transcripts is then passed to subsequent library preparation steps.

#### Transient transcriptome (TT)-seq

The TT-seq experimental mode simulates a TT-seq style experiment by performing *in silico* fragmentation (see below) directly after full-length RNA generation and labeling but before nascent RNA enrichment. After this fragmentation, *SPARK* enriches for nascent RNA fragments (rather than nascent RNA molecules). Fragments that do not contain at least one nucleotide analog are discarded from the population of fragments before proceeding to library preparation.

#### Precision Nuclear Run-on (PRO)-seq

The PRO-seq experimental mode simulates a run-on assay by simulating the incorporation of a biotinylated nucleotide (biotin-NTP, where N can refer to A|C|G|U and can be user-defined) during the *in silico* transcription process. Users specify an incorporation rate, which defines the probability of a biotin-NTP being incorporated instead of a canonical nucleotide. The upstream most biotin-NTP incorporation event then defines the 3’ terminus of the nascent transcript by truncating the molecule at that nucleotide. Truncated molecules then proceed to the *in silico* fragmentation module (see below), in which only the 3’-most fragment containing the singular biotin-NTP is retained for subsequent library preparation simulation.

#### Native Elongating Transcript (NET)-seq

The NET-seq experimental mode simulates RNAP pulldown-based nascent RNA sequencing assays by identifying all molecules actively undergoing elongation at the end of the simulated experiment time. The simulation mimics RNAP pulldown followed by MNase digestion. For each actively transcribing molecule (*i.e.* RNAP has not reached the 3’ end of the gene), *SPARK* extracts the 3’-terminal fragment that represents the nascent RNA physically protected by the RNAPII complex by selecting the last N nt of the molecule, where N is drawn from a uniform distribution between 35-100 nucleotides^4^. Only these 3’ fragments are retained for subsequent library preparation steps.

#### Chromatin-associated RNA enrichment

The chromatin-associated RNA experimental mode simulates biochemical fractionation of the chromatin-bound RNA pool. *SPARK* partitions the simulated molecules into three classes based on their 3′ end processing state: transcripts still undergoing elongation whose 3′ ends have not been cleaved, the nascent readthrough fragments downstream of the CS that remain tethered to the elongating polymerase, and the cleaved molecules. Because uncleaved transcripts and readthrough fragments are, by definition, still engaged with RNAP, *SPARK* retains them in the chromatin fraction. The cleaved RNAs, which are no longer physically anchored to RNAP, are subject to a stochastic, first-order release process. Users define release kinetics by providing a global range from which the framework samples a single gene-specific chromatin-release half-life, representing the time required for half of the cleaved transcript population to dissociate from chromatin. For each molecule, the probability that it remains chromatin-associated decays exponentially with the time elapsed since its cleavage. The retained cleaved molecules, together with all uncleaved transcripts and readthrough fragments, constitute the enriched chromatin-associated population that is passed to subsequent library preparation steps.

### *SPARK in silico* library preparation

The final component of *SPARK* is to perform *in silico* library preparation from the enriched nascent RNAs following simulation of experimental approaches, resulting in high-throughput sequencing reads.

#### Fragmentation and size selection

For experimental modes that include a random fragmentation step during library preparation (e.g., 4sU-, TT-, and PRO-seq), *SPARK* simulates RNA fragmentation as described before^21,31^. Briefly, the length of the fragments is drawn from a modified Weibull distribution with δ = *log*_10_ (*lengt*ℎ*_mRNA_*) and η = mean of the selected fragmentation length interval. This will sample *N_transcripts_* such that:

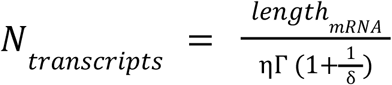

These breakpoints then give *n* length that can be transformed into a fragment of length (*di*) by:

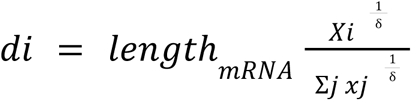

For experimental modes that involve metabolic labeling (nascent RNA pulldown and TT-seq), only fragments within the specified fragment length range are retained in the final fragment pool. Neither PRO- nor mNET-seq approaches have size selection steps.

#### Read generation

*SPARK* can generate either short or long high-throughput sequencing reads. For short reads, simulated reads are generated from *N* nucleotides at the beginning, end, or both (for paired-end sequencing) of the fragments or full-length molecules if no fragmentation was performed), where *N* is a user-specified read length. The default library orientation simulates a dUTP-based protocol (fr-firststrand). For a paired-end simulation, this results in the first read (R1) being the reverse complement of the RNA strand, and the second read (R2) having the same orientation as the RNA strand. *SPARK* can also simulate fr-secondstrand and unstranded library configurations (**Fig. S2B**). For the generation of long reads, the entire simulated mRNA molecule is output as a read. Users can specify the strand orientation to emulate different long-read technologies. This includes options for 3’ to 5’ strandness (mimicking direct RNA sequencing) or unstranded reads (5’ to 3’ or reverse complement 3’ to 5’, mimicking cDNA sequencing), as seen in Oxford Nanopore library preparation methods.

#### Sequencing Errors

*SPARK* can simulate stochastic sequencing errors. A user-specified rate dictates the probability that any given nucleotide in a simulated read will be substituted with a different, randomly chosen nucleotide. The default error rate is 0.01%, reflecting the typical error profile of Illumina sequencing platforms. For long-read sequencing, this error rate should likely be raised to 5-10%.

### *SPARK* optional experimental conditions

#### Background molecules

Since most nascent RNA genomic approaches involving either biotin or antibody-based pulldowns contain some level of background pulldown, *SPARK* allows users to include background molecules from the simulated genes by specifying the proportion of contaminating molecules coming from noisy pulldowns.

#### Nucleotide recoding

The usage of metabolic labeling with synthetic nucleotide analogs allows for nucleoside recoding, in which substitutions are introduced during reverse transcription and library preparation (e.g., T>C or G>A for 4-thiouridine and 6-thioguanine, respectively) at locations of an incorporated analog. To simulate nucleotide recoding, *SPARK* allows users to specify the desired substitutions and a re-encoding rate, which defines the probability that a substitution will be introduced at any given position where a nucleotide analog has been incorporated (which is conditional on the incorporation rate above).

#### RNA synchronization

*SPARK* can simulate the synchronization of RNAP transcription, in which drugs (e.g. DRB) are used to block new initiation or elongation for a time period that allows ongoing elongation to complete RNA synthesis before allowing new transcriptional cycles to start. This is a common experimental strategy (e.g., 4sUDRB-seq) to measure elongation rates. In this mode, the simulation does not generate a steady-state population of pre-existing, elongating polymerases. Instead, all simulated nascent transcripts originate from RNAP molecules that initiate transcription after the simulated experimental start time, mimicking a synchronized release from the TSS. The initiation of these new transcripts on each gene copy still follows the stochastic Poisson process described previously.

## 4sUDRB seq replication

To replicate empirical 4sUDRB-seq data, we obtained published elongation rates for 1,474 human genes from Fuchs *et al.* 2014^18^. Using these rates, we simulated 4sUDRB-seq reads in *SPARK* using read lengths and 4sU labeling duration (8 minutes) matching the experimental data from which the rates were derived. We aligned both *SPARK*-simulated and empirical sequencing reads with STAR^32^ and subsampled aligned reads to obtain the same total read counts per gene between simulated and experimental datasets. We evaluated coverage concordance between datasets by calculating Pearson correlation coefficients across non-overlapping genomic bin sizes of 250, 500, and 1000 nt. To eliminate confounding signals driven by differences in PPP duration, we excluded the initial 1,500 nt of the gene.

### PRO-seq replication

To replicate empirical PRO-seq data, we obtained published TSS pausing durations for 4,481 human genes from Zhang *et al.* 2025 ^22^. Using these rates, we simulated PRO-seq reads in *SPARK* using read lengths and nuclear run-on duration (8 minutes) matching the experimental data from which the pause durations were derived. We aligned both *SPARK*-simulated and empirical sequencing reads with STAR^32^. We evaluated coverage concordance between datasets by calculating Pearson correlation coefficients across non-overlapping genomic bin sizes of 250, 500, and 1000 nt within the first 2,500 nt of the gene.

## 4sU replication

To replicate empirical 4sU-seq data, we obtained published intron half lives for 17,670 introns across 5,475 *Drosophila melanogaster* genes. Using these rates, we simulated 4sU-seq reads in *SPARK* using read lengths and 4sU labeling time (5 minutes) matching the experimental data from which the rates were derived. We aligned both *SPARK*-simulated and empirical sequencing reads using STAR^32^. We calculated the proportion of intron-exon and exon-exon junction reads in each dataset and correlated the empirical and estimated values.

### Parameters used for library quality control

For this manuscript, all simulated datasets were generated using a strandedness setting that emulates dUTP-based library preparation (fr-firststrand), simulated read lengths of 2 x 100nt reads, and library insert sizes of 200 - 300nt, unless specified otherwise.

#### Gene expression estimation

We simulated 3,000 fully synthesized and processed genes to emulate a polyA-selected mature RNA-seq library. TPMs were calculated from simulated fastq files using kallisto^33^.

#### Fragment size estimation

Simulations were performed for 3,000 genes using the metabolic labeling mode across a range of insert size distributions. Following read mapping, insert size distributions for each library were estimated using the CollectInsertSizeMetrics command from Picard^34^.

#### Sequencing error estimation

Simulations were performed for 500 genes using the metabolic labeling mode. Simulated reads were aligned to the hg38 reference genome using STAR^32^ with the following parameters: --outSJfilterOverhangMin 20 12 12 12, --outSJfilterCountUniqueMin 2 1 1 1, --alignIntronMin 25, and --alignIntronMax 250000 and the percentage of unmapped reads was obtained from STAR’s summary.

### FLAM-seq benchmarking

To benchmark the FLAM-seq analytical pipeline^24^, we used SPARK to simulate a long-read sequencing dataset for mature RNAs across 200 genes, assigning variable poly(A) tail lengths ranging between 10 and 200 As per transcript. To reproduce the library structure from FLAM-seq, we added a stretch of Gs to the 3’ end of each simulated transcript. We then processed the resulting reads through the complete FLAM-seq computational pipeline, aligning the long reads to the reference genome using STARlong and estimating per-transcript poly(A) tail lengths.

### GRAND-SLAM benchmarking

To benchmark the GRAND-SLAM pipeline^10^, we used SPARK to simulate metabolic labeling experiments (equivalent to eSLAM-seq) containing T-to-C conversions across varying proportions of unconverted background molecules (mature mRNAs), which represent pre-existing RNA transcripts. For each background proportion, we simulated the same 1,000 genes and generated 100 nt paired-end reads. We aligned the simulated reads to the reference genome using STAR^32^ and used the resulting alignment files in GRAND-SLAM to estimate gene-specific new to total ratios (NTRs) per gene.

### Edge effect correction

We use *SPARK* to generate a dataset with which we could develop a new method to correct for edge effects created during short-read library preparation size selection steps. Simulations were performed for 50 human genes across gene expression levels or fragment size ranges using the nascent RNA pulldown mode. All other biological and technical parameters were held constant. After mapping, base-specific coverage was obtained with bedtools coverage. Though edge effects occur at both ends of RNA molecules, we focused our analyses on 5’ regions most affected by this edge effect. To do so, we consider the region from the transcription start site to position *x_end_* = 2*f_min_* , where *f_min_* denotes the minimum fragment length selected during size selection. This window size proves to be sufficient to capture the full extent of the 5’ edge effect in our simulations.

We developed a probabilistic correction for the coverage data at the previously defined 5’ region that models the probability of a given nucleotide being affected by the edge effect. Rather than exactly replicating the simulator’s internal fragmentation mechanics, we use approximations that simplify the math and yield a tractable correction. The core idea is to approximate a “dropping probability” *P_drop_*(*x*) that captures the probability of a read being dropped at any position *x*.

Using this quantity, we can compute a corrected coverage level:

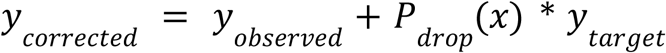

where

*y_observed_*denotes the observed coverage (smoothed using a Savitsky-Golay filter of order 3) and *y_target_* denotes an estimated level of coverage that would have been achieved if no size selection had been performed. Empirically, we set:

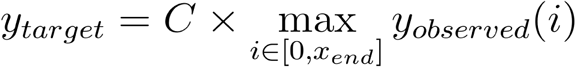

Here, we use the maximum observed coverage in the defined region as a conservative proxy for the target coverage, with *C* being a user-chosen scaling constant that allows for adjustments for global coverage level differences. Throughout our analyses, we set *C* = 0. 75.

Due to constraints from the transcript boundary, the upstream-most fragment of the transcript has a different length distribution than interior fragments, which must be incorporated into the approximation of *P_drop_*(*x*). Since we are primarily interested in the effect of dropping short fragments at this edge, we assume that the internal fragment lengths follow a Weibull distribution with scale parameter *t_min_*denoting the minimum fragment length after size selection and shape parameter *log*_10_(*L*) where *L* is the length of the longest transcript observed. By symmetry, we assume the upstream-most fragment length *d*_1_ also follows a Weibull distribution with identical shape but half the scale. Using this framework, the probability that a position *x* is included in the upstream-most fragment and dropped is given by:

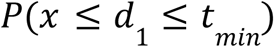

The probability expression can be calculated as *F_d_*_1_ (*t_min_*) − *F_d_*_1_ (*x_d_*_1_), where *F* denotes the cumulative distribution function of *d*_1_ . Since the scale parameter that governs *d*_1_ is halved, the probability of *d*_1_ exceeding the maximum fragment length cutoff *t_max_* is empirically very close to zero and not included in the correction above.

Coverage depletion in the 5’ edge region does not arise solely from the upstream-most fragment. We therefore also consider the effect of the adjacent fragment immediately downstream (the second upstream-most fragment). We are interested in the probability:

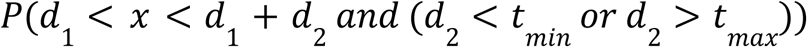

which denotes the probability that the position *x* is covered by a second fragment of length *d*_2_ that is dropped. Since *d*_2_ is the length of an internal fragment, it is assumed to be independent of *d*_1_ and follows the same Weibull distribution that governs the length of internal fragments. Since the distributions of both *d*_1_ and *d*_2_ are defined exactly, this probability can be accurately and efficiently evaluated in Python using a standard numerical integration/quadrature function from the library SciPy. We can therefore approximate *P_drop_*(*x*) as:

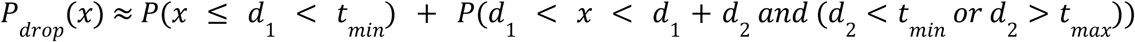

This quantity can then be used to estimate *y_corrected_*.

To evaluate the efficacy of our probabilistic correction, we compared it against both the uncorrected raw data and a baseline linear corrector. The linear corrector is computed by fitting a simple linear regression model against coverage data in the region from the transcription start site to 2 × *x_end_* . We compared the coverage levels yielded from these three approaches (raw, linear, and the probabilistic corrector) against the ground truth coverage using the symmetric mean absolute percentage error (symmetric MAPE). More precisely, given the ground truth coverage level *y_trut_*_ℎ_ and the corrected coverage level *y_corrected_*, the symmetric MAPE is given by:

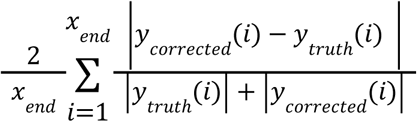

For each fragment size or gene expression level and correction method, we computed the symmetric MAPE for all 50 genes and performed pairwise t-tests to gauge the statistical significance of any performance differences after Bonferroni correction for multiple-hypothesis testing.

