## Supplementary Material for "A mechanistic framework for nascent RNA-seq simulations"

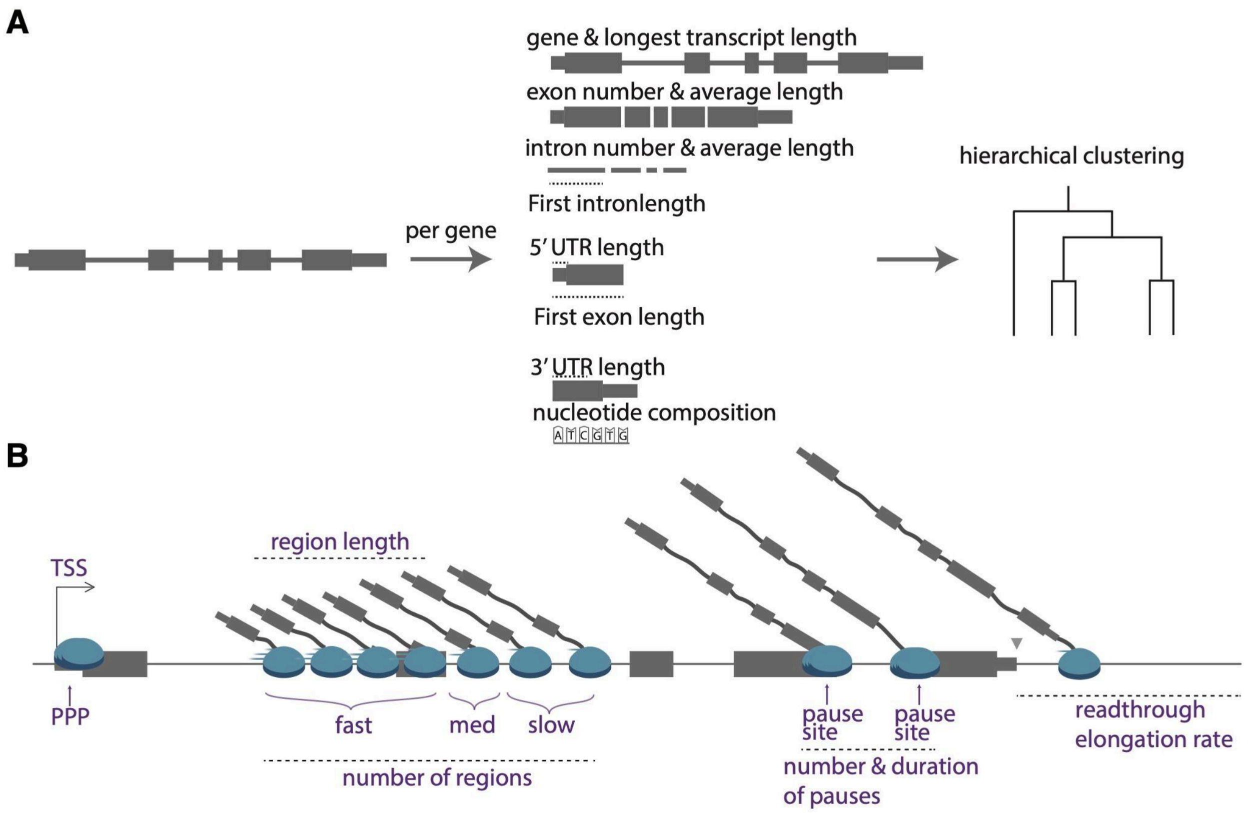


**Supplementary Figure 1. Gene selection and user-defined RNAP elongation parameters for *SPARK*. (A)** Genes are categorized by features such as exon or intron count, total exonic or intronic real estate, and 5' or 3' UTR lengths. These features are used as inputs for hierarchical clustering to select a diversity of genes from which to simulate transcripts and reads. Users can define the number of genes to simulate and the number of clusters across which to distribute genes. **(B)** Biological and technical parameters simulated by *SPARK*. In addition to ranges of transcription initiation and RNAP elongation rates, users can define promoter proximal pausing duration, the number and length of regions over which RNAP elongation rates vary, and the number and duration of gene-body pause sites.

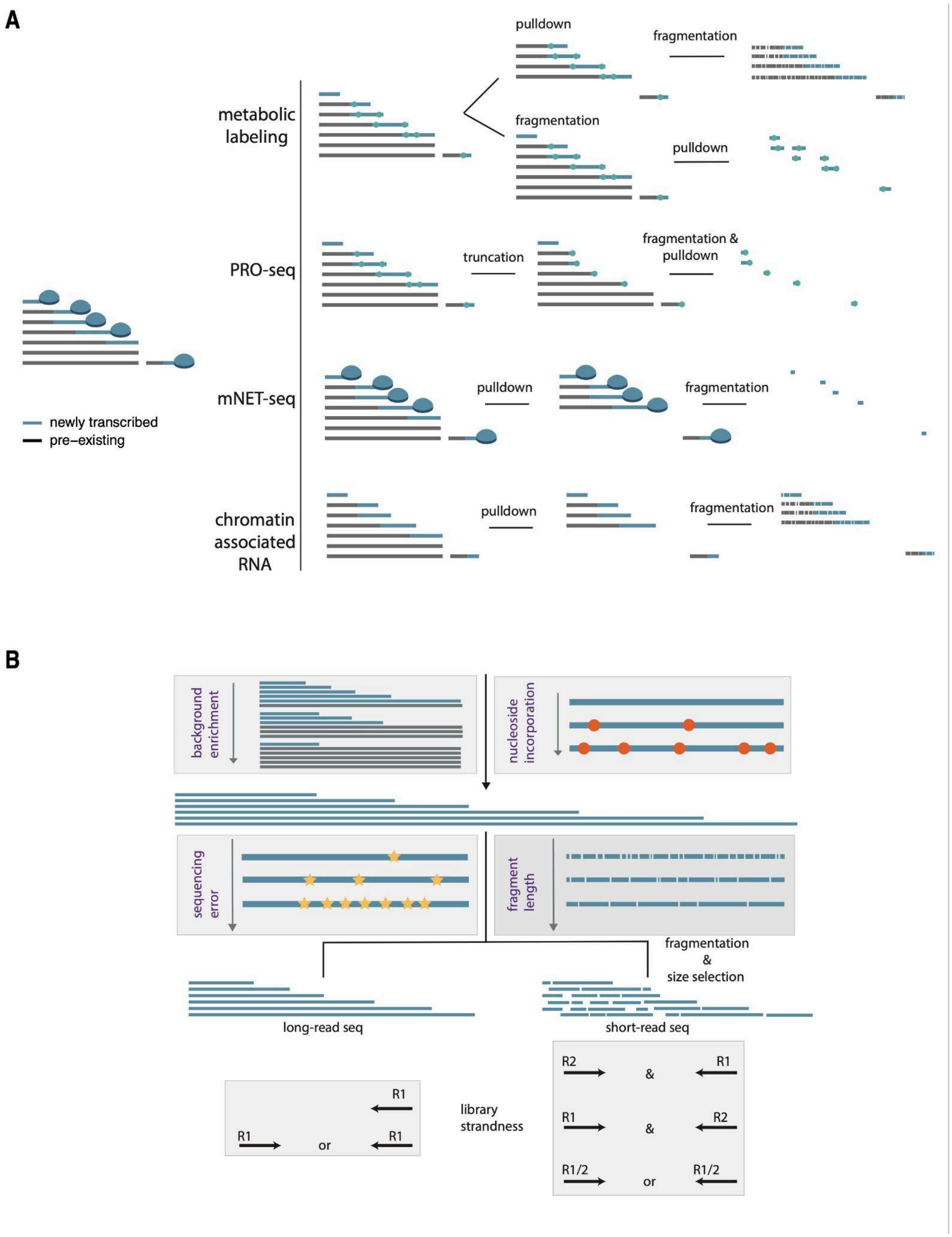


**Supplementary Figure 2. Nascent RNA enrichment methods and user-defined parameters for *SPARK*. (A)** Schematic representation of the nascent RNA enrichment and library preparation workflows modeled in *SPARK* for metabolic labeling. Blue bars represent newly transcribed RNA; grey bars represent pre-existing RNA. **(B)** Users can define the frequency at which *SPARK* introduces background (non-nascent) molecules and re-coded nucleoside-derived substitutions. The library preparation module simulates fragmentation (if appropriate and at the correct stage in the protocol), size selection of fragments for either long-read or short-read sequencing, library strandness, and random sequencing errors, all of which can be tuned by users.

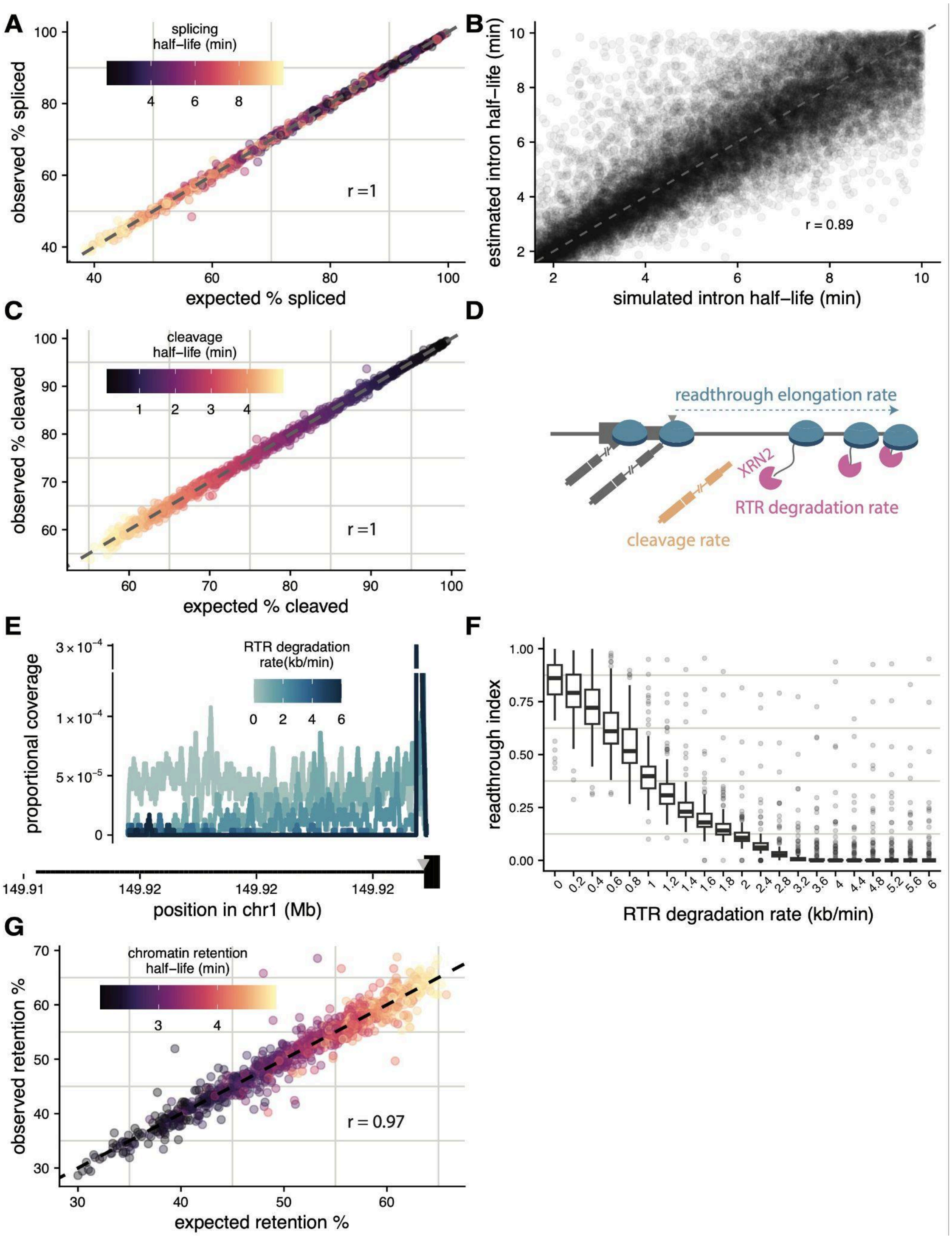


**Supplementary Figure 3. Simulated RNA processing kinetics. (A)** The correlation between expected (*x-axis*) and observed (*y-axis*) percentages of spliced introns across simulated half-lives (n = 1000, Pearson's r = 1). The grey line represents y = x. (**B**) Genome-wide correlation between ground truth simulated intron half-lives (*x-axis*) and estimated half-lives (*y-axis*)across 35,000 introns from 4,000 genes (Pearson's r = 0.89). **(C)** The correlation between expected (*x-axis*) and observed (*y-axis*) percentages of cleaved transcripts (n = 1000, Pearson's r = 1). The grey line represents y = x. **(D)** Schematic overview of RNA processing dynamics at the 3' end of a transcript, including the rate of cleavage at the cleavage site (*grey triangle*), the rate of RNAP elongation in the readthrough region, and the degradation of the readthrough transcript (presumably by XRN2 digestion). **(E)** Read coverage track for the gene *SF3B4* (reverse strand) across varying readthrough degradation rates. The grey triangle indicates the cleavage site. **(F)** Genome-wide distribution of the Readthrough Index (RTI; *y-axis*) as a function of the readthrough degradation rate (*x-axis*, n = 200 per degradation bin). **(G)** Correlation between expected (x-axis) and observed (*y-axis*) chromatin retention percentages across simulated chromatin retention half-lives (r = 0.97). The grey line represents y = x.

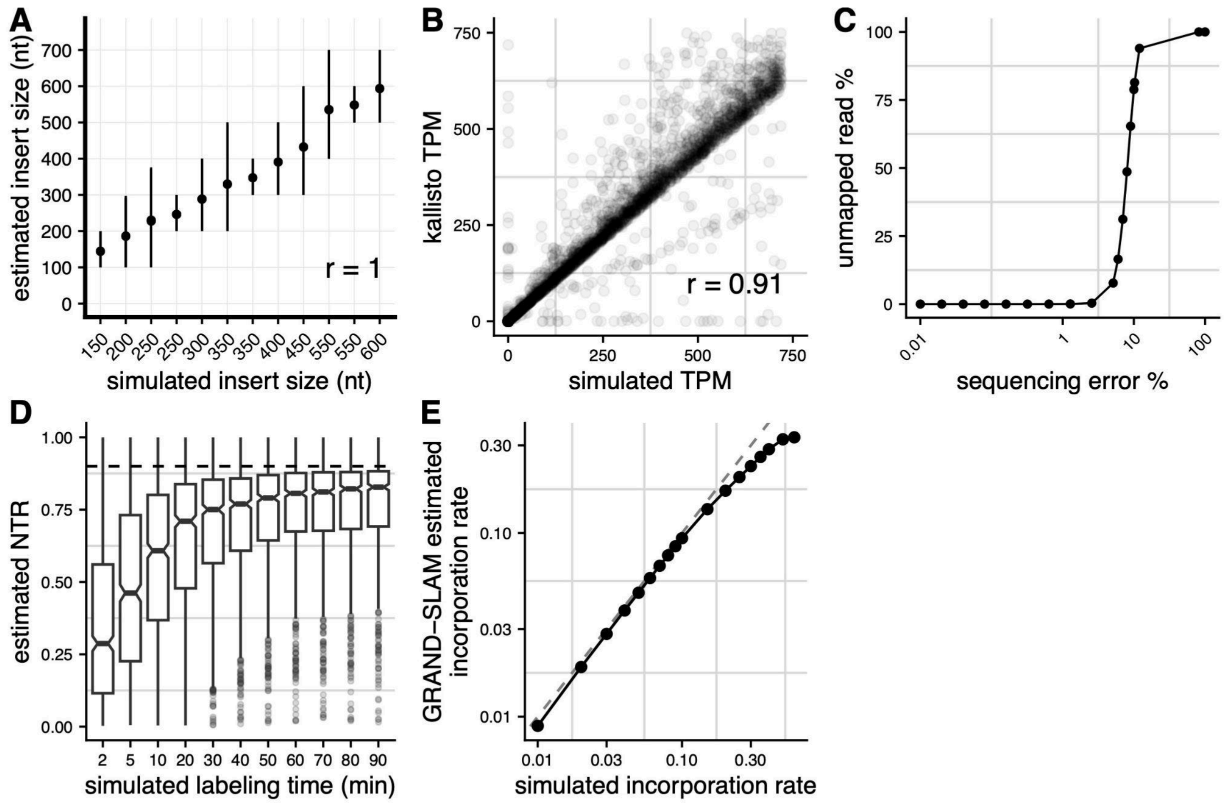


**Supplementary Figure 4. Validation of library preparation parameters.** Correlations between simulated parameters (*x-axis*) and estimated parameters from simulated reads (*y-axis*) for **(A)** library insert sizes (Pearson's r = 1, where points represent mean and vertical bars show full range of detected fragments) and **(B)** gene expression levels. **(C)** The percentage of unmapped reads (*y-axis*) as a function of the simulated sequencing error rate (*x-axis*). (**D**) The distribution of GRAND-SLAM estimated NTRs (*y-axis*) across simulated 4sU labeling times (*x-axis*) ranging from 2 to 90 minutes at a fixed 4sU incorporation rate of 0.04. The dashed horizontal line represents the expected *in silico* ground-truth NTR of 0.9. (**E**) Correlation between the ground-truth simulated 4sU incorporation rate (*x-axis*) and the incorporation rate estimated by GRAND-SLAM (*y-axis*). The dashed diagonal line represents y=x.
